# Human embryoid bodies model basal lamina assembly and muscular dystrophy

**DOI:** 10.1101/765131

**Authors:** Alec R. Nickolls, Michelle M. Lee, Kristen Zukosky, Barbara S. Mallon, Carsten G. Bönnemann

## Abstract

The basal lamina is a specialized sheet of dense extracellular matrix (ECM), linked to the plasma membrane of specific cell types in their tissue context, that serves as a structural scaffold for organ genesis and maintenance. Disruption of the basal lamina and its functions is central to many disease processes, including cancer metastasis, kidney disease, eye disease, muscular dystrophies, and specific types of brain malformation. The latter three pathologies occur in the dystroglycanopathies, which are caused by dysfunction of the ECM receptor dystroglycan. However, opportunities to study the basal lamina in various human disease tissues are restricted due to its limited accessibility. Here, we report the generation of embryoid bodies from human induced pluripotent stem cells to model basal lamina formation. Embryoid bodies cultured via this protocol mimic pre-gastrulation embryonic development, consisting of an epithelial core surrounded by a basal lamina and a peripheral layer of ECM-secreting endoderm. In dystroglycanopathy patient embryoid bodies, electron and fluorescence microscopy revealed ultrastructural basal lamina defects and reduced ECM assembly. By starting from patient-derived cells, these results establish a method for the *in vitro* synthesis of patient-specific basal lamina and recapitulate disease-relevant ECM defects seen in muscular dystrophies. Finally, we applied this system to evaluate an experimental ribitol supplement therapy on genetically diverse dystroglycanopathy patient samples.

## Introduction

Metazoan life relies on tissue compartmentalization to form ordered, discrete organs. This is partly accomplished by an extracellular matrix (ECM) barrier called the basal lamina, which ensheaths epithelial, endothelial, adipose, muscle, and nervous tissue [36]. The main components comprising the basal lamina are laminin isoforms, perlecan, nidogen, and collagen type IV, forming a complex lattice anchored to cell surface receptors [51]. This cell-ensheathing basal lamina is generally inter-connected, on its acellular matrix side, to a “lamina reticularis” composed of fibrillar collagens, microfibrils, and proteoglycans. Together, they form a multilayered basement membrane with tissue-specific mechanical properties [41, 51]. The terms “basal lamina” and “basement membrane” are sometimes used interchangeably. However, here they will refer to distinct structures, with the basal lamina being the dense, cell-attached component of the basement membrane.

The basal lamina is an essential structural component for organ genesis and maintenance. Its perturbation is linked to many human clinical conditions including metastatic cancer, nephropathy, lissencephaly, and muscular dystrophy. One mechanistic group of basal lamina-related diseases pertains to the dysfunction of the cell membrane ECM receptors integrin and dystroglycan [15, 47]. These receptors mediate cell attachment to the basal lamina, and in turn they influence the arrangement of the basal lamina itself [20, 30, 31, 35].

There is an expanding literature on the spectrum of disorders caused by dystroglycan receptor dysfunction, collectively termed the dystroglycanopathies, which are estimated to constitute roughly a third of all congenital muscular dystrophies [17]. A hallmark of severe dystroglycanopathies is rupture or detachment of the basal lamina that encases the brain and muscle fibers during development and structural maintenance [10, 25]. This specific combination of basal lamina abnormalities is associated with a range of developmental nervous system malformations and progressive skeletal muscle degeneration that may ultimately be fatal [38].

The biochemical basis of the dystroglycanopathies is a reduction in a highly specific form of O-linked glycosylation on dystroglycan’s α-subunit (αDG). This leads to a “hypoglycosylation” of the final αDG glycoepitope, which is referred to as the matriglycan [49]. Matriglycans on αDG confer binding activity to the ECM molecules laminin, perlecan, and nidogen [4]. Hypoglycosylated matriglycans have limited ECM binding capacity, which is thought to destabilize the basal lamina in muscle and brain tissue as a common disease pathway in the dystroglycanopathies [34, 35]. The 17 genes that are known to be mutated in the dystroglycanopathies all affect the formation of matriglycans and include various specific glycosyltransferases as well as enzymes preparing specific sugars to be incorporated, while only very few cases involve the gene encoding dystroglycan itself. Based on this knowledge, a large proportion of the dystroglycanopathy cases can now be clarified genetically [8, 17].

Understanding the mechanisms of pathogenesis and developing rational therapies for the dystroglycanopathies remains a challenge, in part due to its phenotypic and genetic heterogeneity. A large collection of dystroglycanopathy animal models recapitulates many aspects of the clinical spectrum [38]. However, such approaches fall short of modeling the genetic diversity of human patients for assessing disease phenotypes and drug responses.

To study patient-specific basal lamina in a model system, we developed a protocol to generate ECM-containing spheroids from human induced pluripotent stem cells (hiPSCs), which we refer to as embryoid bodies. hiPSC-derived embryoid bodies produce their own basal lamina and represent a simplified 3D system to investigate human ECM and its receptors in diverse genetic contexts. As a proof of concept, we applied this method to produce embryoid bodies from a variety of dystroglycanopathy patients. We observed subtle basal lamina defects that correlated with disease severity and corroborate findings in mouse models. Lastly, we evaluated patient hiPSCs and embryoid bodies treated with the sugar alcohol ribitol, a recently proposed therapeutic for the dystroglycanopathies. By correlating patient genotype and drug response, this approach allows for pre-clinical prediction of therapeutic efficacy in specific individuals.

## Materials and methods

### Generation of hiPSC Lines

Written informed consent for patient participation was obtained by a qualified investigator (protocol 12-N-0095 approved by the National Institute of Neurological Disorders and Stroke, National Institutes of Health). Dystroglycanopathy patient hiPSCs were reprogrammed from dermal fibroblasts using an hOKSML mRNA reprogramming kit (Stemgent, 00-0067). Control-1 hiPSCs were reprogrammed in the same manner from BJ foreskin fibroblasts (ATCC, CRL-2522). Immunocytochemical validation of germ layer differentiation was performed off-site (Stemgent). Control-2 hiPSCs were reprogrammed from control foreskin fibroblasts (ATCC, CRL-2097) using the CytoTune-iPS 2.0 Sendai reprogramming kit (Thermo Fisher, A16517). Control-3 hiPSCs (NC15) were previously generated by lentiviral reprogramming of adult dermal fibroblasts [18]. Karyotype analysis was performed after at least 10 passages (WiCell), and all cell lines were routinely tested for mycoplasma contamination (LT07-118, Lonza).

### hiPSC Culture

Human hiPSCs were maintained with daily changes of E8 medium (Thermo Fisher, A1517001) on tissue culture-treated polystyrene plates coated with Matrigel (Corning, 354277) and passaged every 4 – 6 days using ReLeSR (STEMCELL Technologies, 05872).

### Embryoid Body Differentiation

Differentiation of human hiPSC-derived embryoid bodies was performed essentially as described previously for human embryonic stem cells [45]. At least one passage before differentiation, hiPSCs were transitioned to MEF co-culture. The MEFs (Millipore, PMEF-CF) were seeded at 30,000 cells/cm^2^ on plates coated with gelatin (STEMCELL Technologies, 07903) and maintained in serum-containing medium consisting of KO-DMEM (10829-018), 20% FBS (26140-079), 100 µM non-essential amino acids (11140-050), 2 mM GlutaMAX (35050-061), and 55 µM β-mercaptoethanol (21985-023) (all from Invitrogen).

hiPSCs were dissociated with ReLeSR and plated on MEFs in knockout serum replacement medium consisting of KO-DMEM, 20% KSR (Invitrogen, 10828-028), 100 µM non-essential amino acids, 2 mM GlutaMAX, 55 µM β-mercaptoethanol, 10 ng/mL bFGF (Thermo Fisher, 233-FB-025), and 10 µM Y-27632 (Tocris, 1254). The medium was changed daily (without Y-27632) until hiPSCs reached roughly 60% confluency.

For embryoid body formation, hiPSCs were dissociated by Collagenase Type IV (Invitrogen 17104-019) and manual scraping followed by gravity sedimentation to remove as many MEFs as possible. The cells were then individualized with Accutase (Invitrogen, A1110501), and 2.4 x 10^6^ cells were seeded per well of an AggreWell 400 (STEMCELL Technologies, 34411) following manufacturer’s instructions by centrifugation in X-VIVO 10 medium (Lonza, 04-380Q) with 10 µM Y-27632. The following day, spheroids were extracted from the AggreWell 400 according to manufacturer’s instructions and cultured in ultra-low attachment dishes (Corning, 3262) with serum-containing medium for up to four days with a medium change every other day.

### Endoderm-Free Embryoid Body Culture

Before endoderm-free embryoid body experiments, feeder-free hiPSCs were maintained on Matrigel in E8 medium. hiPSCs were dissociated with Accutase, and 1.2 x 10^6^ cells were seeded per well of an AggreWell 400 by centrifugation in E8 with 10 µM Y-27632. The next day, spheroids were transferred to ultra-low attachment 6-well plates (Corning, 3471) in E8 supplemented with laminin (Invitrogen, 23017015) for 48 hours. 140µg/mL laminin was used except where otherwise stated in the main text.

### Western Blotting

hiPSCs in 100 mm dishes were lysed by 200 µL RIPA buffer with protease and phosphatase inhibitors. 1 mg soluble protein was incubated overnight at 4 °C in 500 µL RIPA with 50 µL agarose-bound wheat germ agglutinin (Vector Labs, AL-1023) to enrich the glycoprotein fraction. The agarose was washed three times with RIPA, and the glycoproteins were eluted by 5-minute incubation at 95 °C in SDS-PAGE loading buffer. Glycoproteins were ran on 4 – 12% Bis-Tris gels and transferred to PVDF membranes.

All blocking steps and antibody incubations were performed in TBST with 5% milk (glyco-αDG and βDG) or 5% donkey serum (core-αDG). The membranes were probed with antibodies against glyco-αDG, core-αDG, or βDG overnight at 4 °C. Labeling was visualized by chemiluminescence with appropriate secondary HRP-conjugated antibodies on a ChemiDoc XRS+ (Bio-Rad). See Table 2 for a list of antibodies used in this study.

**Table 1.** Characteristics of study subjects.

Characteristics of study subjects
| Subject | Age | Gender | Clinical presentation | Diagnosis | Genotype |
| --- | --- | --- | --- | --- | --- |
| **Control-1** | Neonate | Male | N/A | N/A | N/A |
| **Control-2** | Neonate | Male | N/A | N/A | N/A |
| **Control-3** | 51 | Female | N/A | N/A | N/A |
| **LARGE** | 6 | Female | Delayed motor milestones, weakness and general hypotonia, able to walk stairs, mild autistic behavior, mild muscle biopsy, lissencephaly on brain MRI | Congenital muscular dystrophy type 1D | LARGE1: Heterozygous<br>- Exon 7 deletion<br>- Exons 3 – 7 deletion |
| **FKRP** | 3 | Male | Delayed motor milestones, mild weakness, trunk hypotonia, waddling gait, able to run, mild autistic behavior, moderate dystrophy on muscle biopsy, normal basement membrane on TEM | Early onset limb-girdle muscular dystrophy type 2I | FKRP: Heterozygous<br>- c.C826A (p.L276I)<br>- c.G534T (p.W178C) |
| **POMT2** | 4 ½ | Female | Severely delayed motor milestones, hip dysplasia at birth, weakness and general hypotonia, unable to stand, joint contractures, failure to thrive, delayed speech acquisition, white matter hyperintensities on MRI | Muscle-Eye-Brain Disease | POMT2: Homozygous<br>- c.G1057A (p.G353S) |

Age in years; TEM, transmission electron microscopy; MRI, magnetic resonance imaging

**Table 2.** Antibodies for immunofluorescence and western blots.

| Antibody | Dilution | Company | Catalog Number |
| --- | --- | --- | --- |
| **SOX17** | 1:100 | Abcam | ab84990 |
| **OCT4** | 1:200 | Abcam | ab134218 |
| **Laminin** | 1:1,000 (IF)<br>1:5,000 (WB) | Sigma-Aldrich | L9393 |
| **Glyco-αDG**<br>(IIH6C4) | 1:200 (IF)<br>1:1,000 (WB) | Millipore | 05-593 |
| **Core-αDG** | 1:500 | R&D Systems | AF6868 |
| **βDG** | 1:10,000 | GeneTex | GTX124225 |
| **Nestin** | 1:500 | Millipore | ABD69 |
| **SSEA3** | 1:200 | Abcam | ab16286 |
| **SSEA4** | 1:500 | STEMCELL Technologies | 60062 |
| **F-actin**<br>(Phalloidin-647) | 1:100 | Thermo Fisher | A22287 |
| **Perlecan** | 1:100 | Millipore | MAB1948-P |
| **Nidogen** | 1:50 | R&D Systems | MAB2570 |
| **COLIV** | 1:1,000 | Chemicon | MAB1430 |
| **HuNu** | 1:100 | Millipore | MAB1281 |

IF, immunofluorescence; WB, western blot

### Laminin Overlay Assay

PVDF membranes were first blocked with 5% milk in laminin binding buffer (LBB; 140 mM NaCl, 10 mM triethanolamine, 1mM CaCl_2_, 1mM MgCl_2_, 0.05% Tween, pH 7.6) and then incubated with 1 µg/mL laminin in LBB overnight at 4 °C. PVDF membranes were washed and probed with laminin antibodies for 1 hour at room temperature in LBB with 5% milk. The membranes were then washed and probed with an appropriate HRP-conjugated secondary antibody for 1 hour at room temperature in LBB with 5% milk before chemiluminescent imaging.

### Immunofluorescence Microscopy

For immunocytochemistry, cells in chamber slides were fixed for 10 minutes in 4% PFA and then washed with PBS before staining. For immunohistochemistry, embryoid bodies were fixed for 20 minutes in 4% PFA, cryoprotected by overnight incubation with 30% sucrose in PBS, and frozen in optimum cutting temperature (OCT) compound (VWR, 25608-930). OCT blocks were sectioned at 10 µm thickness on a cryostat and mounted on slides for staining.

Slides were blocked in 10% goat serum and 0.1% Triton X-100 for 1 hour at room temperature before primary antibody incubation with 3% goat serum overnight at 4 °C. Secondary antibody labeling was performed at room temperature for 1 hour. Refer to Table 2 for antibody dilutions and catalog numbers. Fluorescent images were captured on a Leica TSC SP5 II confocal microscope or a Nikon Eclipse Ti-E inverted microscope.

### Transmission Electron Microscopy

Embryoid bodies were fixed for 30 minutes at room temperature in 0.1 M cacodylate buffer with 4% glutaraldehyde, pH 7.4. Samples were then coated in agarose, washed with buffer, and incubated for 60 minutes at 4 °C in 0.1 M cacodylate buffer with 1% osmium tetroxide, pH 7.4. The samples were washed and stained *en bloc* overnight at 4 °C in 0.1M acetate buffer with 1% uranyl acetate, pH 5.0. The next day, samples were dehydrated in ethanol and epoxy resin embedded. 70 nm sections were cut and counterstained with lead citrate and uranyl acetate. Micrographs were captured on a JEOL1200EX transmission electron microscope with a digital CCD camera (AMT XR-100, Danvers, MA, USA).

### Image Quantification and Statistics

The image processing program Fiji was used to analyze all western blots and microscopy images. Staining intensity was measured by drawing equal sized regions of interest and measuring the average pixel intensity in each sample. Before statistical measurements, all data were assessed for normality using the Shapiro-Wilk test. If data were not normally distributed, they were analyzed by Kruskal-Wallis test with Dunn’s correction for multiple comparisons (Fig. 5e). If data were normally distributed, an ordinary one-way ANOVA was used and corrected with Tukey’s multiple comparisons test (all other figures). The number of replicates, n, for each analysis are reported in their respective figure legend. Prism 7.0 (GraphPad software) was used to make all statistical tests and graphs.

## Results

### Human Embryoid Bodies Mimic Pre-Gastrulation Development

To establish an hiPSC-based model of basal lamina assembly, we used a microwell plate to generate spheroids of human hiPSCs (Fig. 1a, b). Following transfer of the spheroids into suspension culture, we tested multiple conditions for optimal ECM production. Spheroids grown in a standard knockout serum replacement (KSR) medium formed a cavitated core and differentiated into a Nestin+ neuroectodermal lineage (Fig. 2a). This result could be achieved with either feeder-free hiPSCs or with feeder-dependent hiPSCs, the latter of which were cultured on a feeder layer of mouse embryonic fibroblasts (MEFs) prior to spheroid formation.

**Fig. 1.**
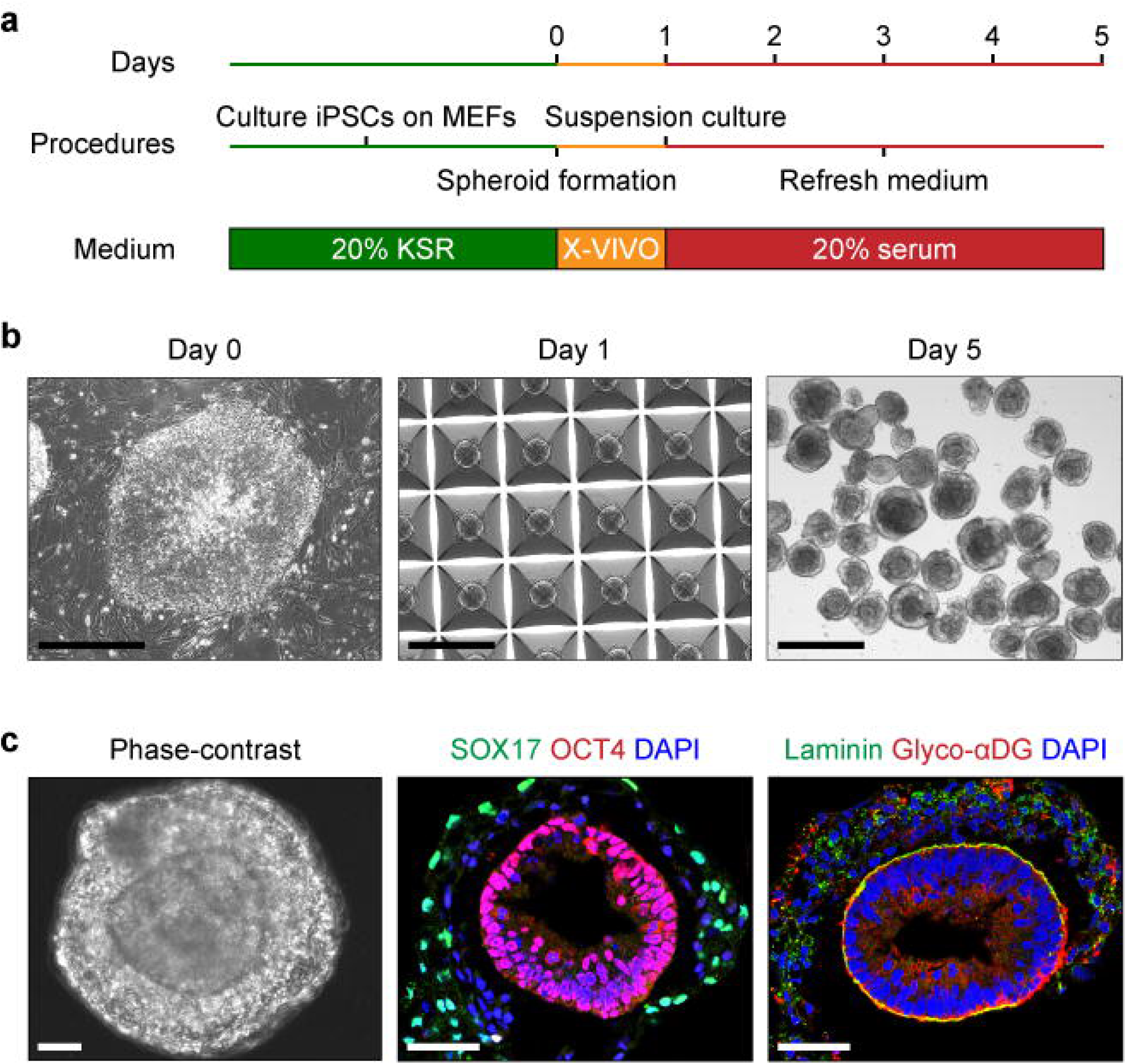
Self-Organization of Basal Lamina-Containing Embryoid Bodies from Human hiPSCs. (**a**) Schematic of embryoid body differentiation protocol from feeder-dependent hiPSCs. X-VIVO refers to X-VIVO 10 medium (see experimental procedures). (**b**) Phase-contrast representation of embryoid body differentiation. hiPSCs were seeded on day 0 in a microwell plate to form spheroids of roughly 2,000 cells on day 1. The spheroids were then maintained in suspension culture until day 5. Scale bars, 500 µm. (**c**) Phase-contrast and immunohistochemistry on day 5 embryoid bodies showing two distinct tissue domains, with a basal lamina in contact with the interior OCT4+ cells expressing glycosylated αDG (glyco-αDG). Scale bars, 50 µm.

**Fig. 2.**
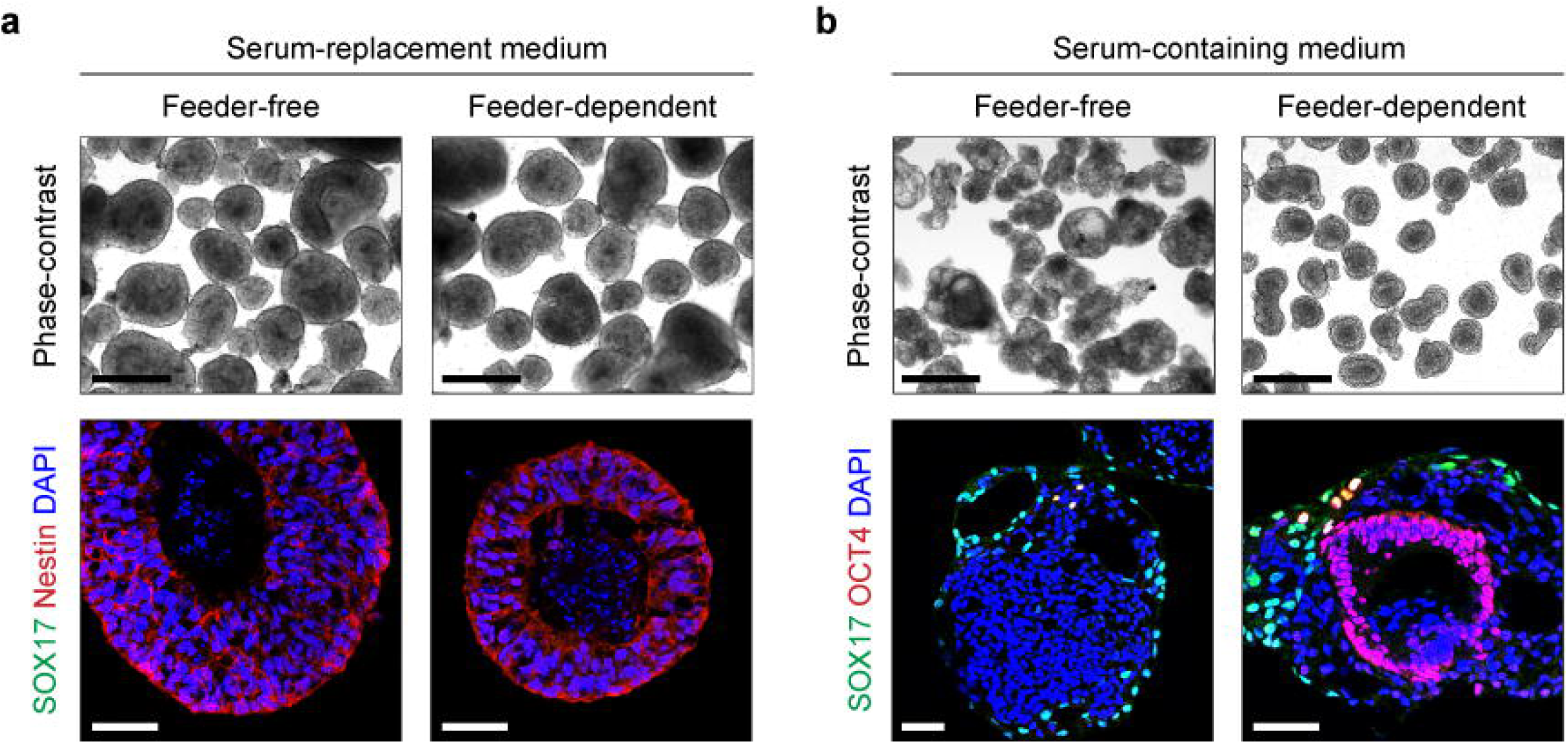
Culture Conditions Impact the Lineage Outcome of Human hiPSC-Derived Embryoid Bodies. (**a**) Phase-contrast and immunohistochemistry of hiPSC spheroids maintained in 20% serum-replacement medium for 5 days. Feeder-dependent spheroids were derived from hiPSCs cultured on a feeder layer of MEFs for at least one passage. (**b**) Phase-contrast and immunohistochemistry of day 5 spheroids in serum-containing medium. Phase-contrast scale bars, 500 µm; fluorescence scale bars, 50 µm.

We next tested the effect of serum on hiPSC spheroid differentiation. In a medium with 20% serum, spheroids from feeder-free hiPSCs consisted of an outer SOX17+ endodermal layer and a disorganized core of differentiated OCT4-cells (Fig. 2b). These spheroids, as well as those discussed above that were maintained in KSR medium, were mostly devoid of ECM as assessed by antibodies against the basal lamina protein laminin (data not shown).

In contrast, we found that feeder-dependent hiPSCs produced spheroids with two visibly partitioned domains when cultured in serum-containing medium (Fig. 1c, 2b). These particular spheroids were characterized by a SOX17+ endodermal periphery and an OCT4+ epithelial core. The SOX17+ cells were often mixed with an additional unidentified OCT4-/SOX17-population. These inner and outer tissue compartments were demarcated by a laminin-rich basal lamina (Fig. 1c).

The SOX17+ endodermal cells were apparently responsible for ECM secretion, with disorganized aggregates of laminin visible in the spheroid outer layer (Fig. 1c). The underlying core of OCT4+ cells appeared as a radially arranged epithelium and expressed the basal lamina receptor αDG, which was enriched at the basal lamina interface between the two tissue domains (Fig. 1c). Surprisingly, as little as one passage on MEFs was sufficient to prime the hiPSCs for this self-organized differentiation. Similar tissue patterning has been reported in spheroids derived from mouse and human embryonic stem cells [30, 45]. The observed structure is thought to represent pre-gastrulation embryonic development, with an epiblast-like core and an outer layer of extra-embryonic endoderm. Because the hiPSC-derived spheroids produced with our protocol resemble this developmental stage, we refer to them hereafter as embryoid bodies.

### Derivation of hiPSCs from Dystroglycanopathy Patients

Because hiPSC-derived embryoid bodies express αDG and produce ECM in the form of a basal lamina, we sought to apply this system for evaluating basal lamina phenotypes in dystroglycanopathy – a diverse spectrum of muscular dystrophies often co-morbid with brain malformation, characterized by ruptures in the basal lamina. We reprogrammed hiPSCs from dermal fibroblasts of three unrelated individuals with a genetic diagnosis of dystroglycanopathy. Each patient harbored predicted pathogenic mutations in a different gene required for the glycosylation of αDG: *LARGE*, *FKRP*, or *POMT2* (Table 1). The LARGE patient has been reported in a previous publication [33].

Clinical presentation for all three patients included delayed motor milestones, muscle weakness, and cognitive impairments (Table 1). Overall, the FKRP patient had the mildest clinical and muscle biopsy findings, while the patient with the POMT2 mutations had the most severe findings. Brain magnetic resonance imaging showed minor white matter and structural abnormalities in the LARGE patient [33] and POMT2 patient (data not shown). Hypoglycosylation of αDG is the primary causative factor in the pathogenesis of dystroglycanopathy [34]. To evaluate the glycosylation status of αDG in patient cells, we used the IIH6C4 antibody that specifically recognizes the glycosylated form of αDG (glyco-αDG). Consistent with clinical and genetic observations, we found reduced expression of glycosylated αDG in our patient hiPSCs (Fig. 3a). Two hiPSC clones were evaluated for each patient. All cell lines had a normal karyotype, and there were no discernible differences in the expression of pluripotent markers or propensity for germ layer differentiation (Fig. 3a, 4a-c).

**Fig. 3.**
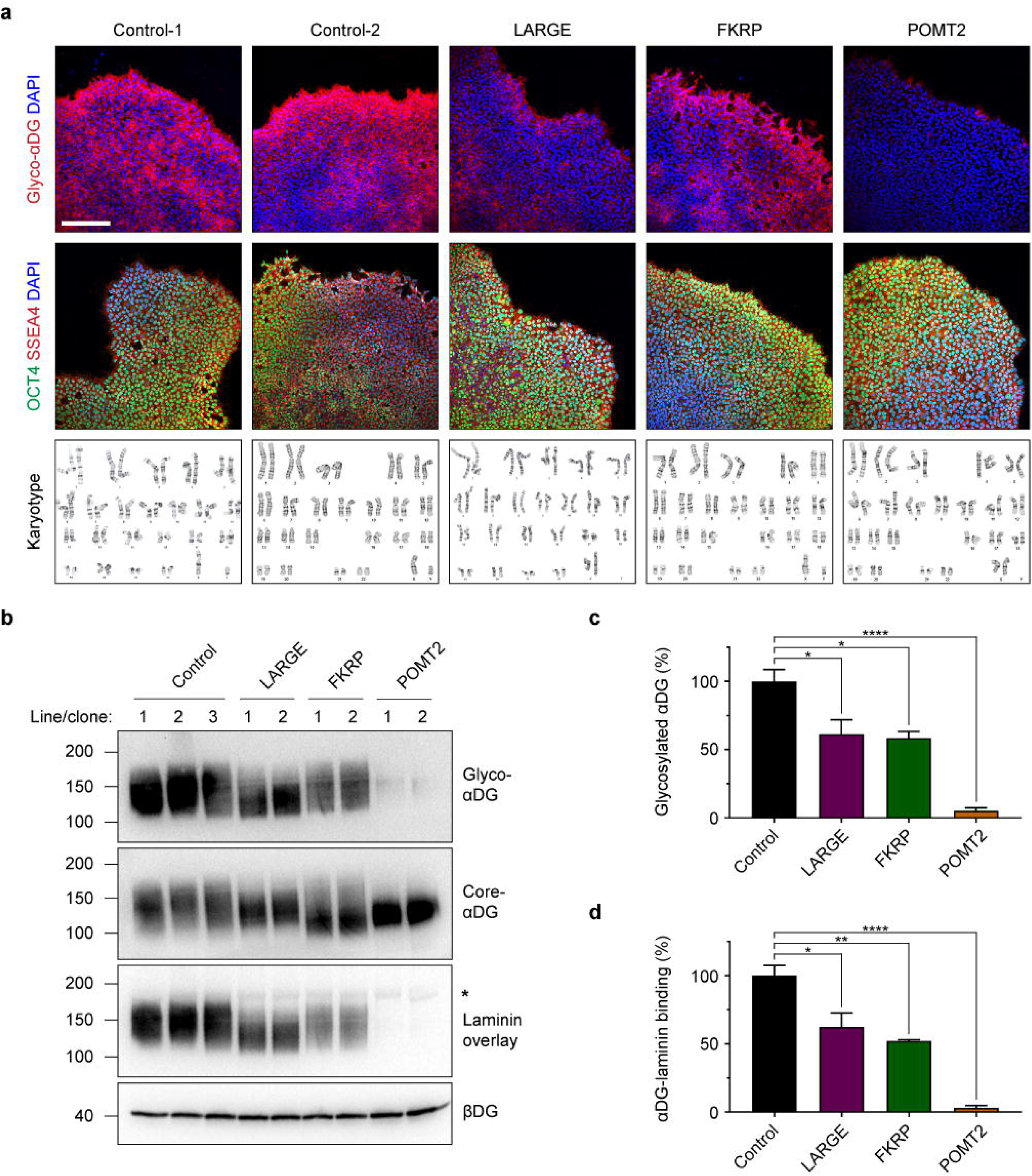
Dystroglycanopathy Patient hiPSCs Express Hypoglycosylated Forms of αDG. (**a**) Immunocytochemistry and karyotype analyses of control and dystroglycanopathy patient-derived hiPSCs. Scale bars, 200 µm. (**b**) Western blots on hiPSC protein lysates. βDG was used as a loading control. The asterisk indicates the molecular weight of endogenous laminin in the samples. Each lane represents one cell line (for controls) or one clone (for patients). (**c**, **d**) Quantification of western blots on glycosylated αDG and the laminin overlay assay. Band intensity for each sample was normalized to βDG, and all samples are expressed as a percent of control. Three control cell lines and two clones per patient were used, n = 3 technical replicates per clone. Values expressed as mean ± s.e.m. Post-hoc comparisons * P < 0.05, ** P < 0.01, **** P < 0.0001.

**Fig. 4.**
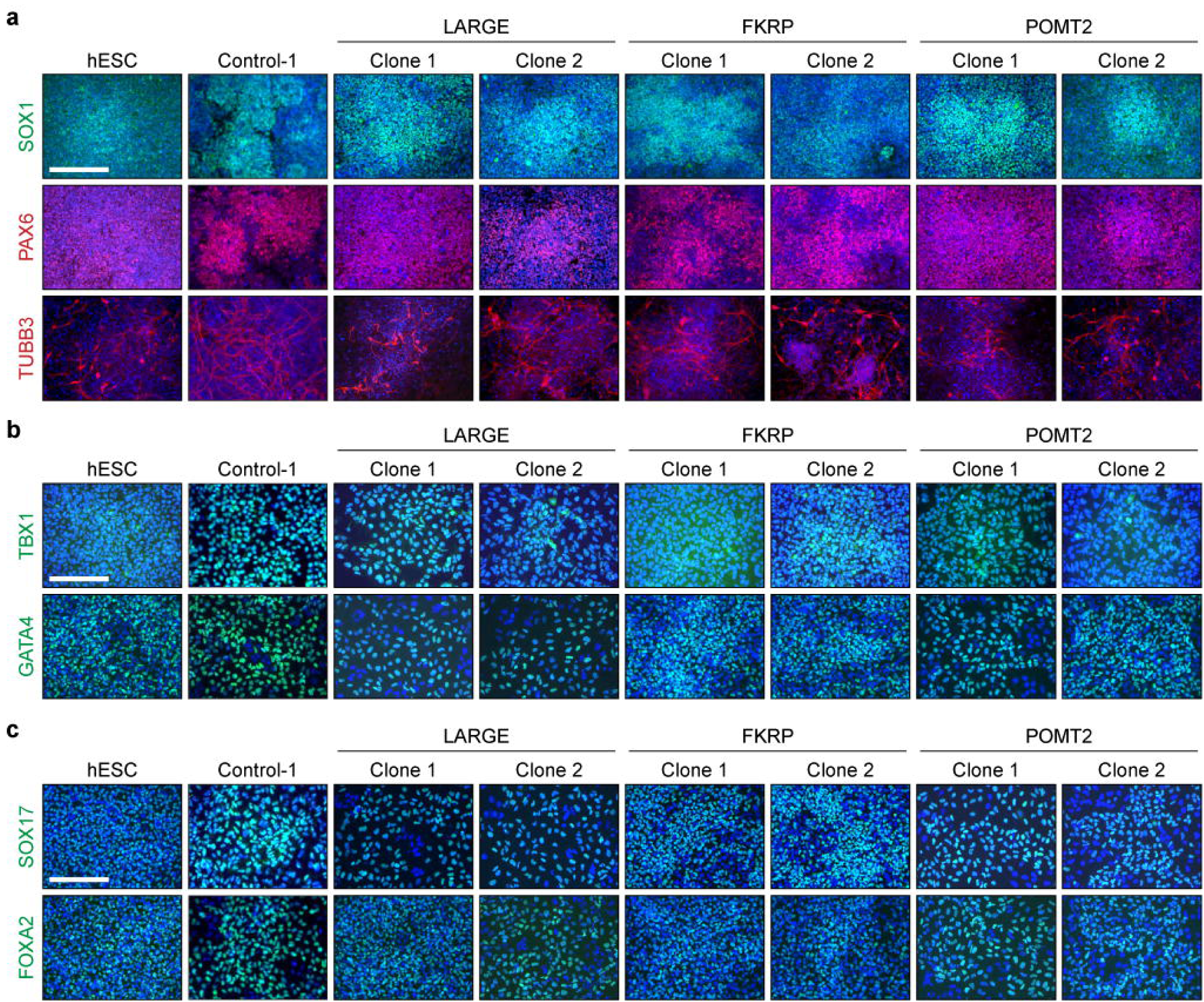
hiPSCs from Dystroglycanopathy Patients Show Normal Germ Layer Differentiation. (**a**) Immunocytochemistry of ectoderm markers. hiPSCs were differentiated via dual SMAD inhibition by treating with LDN-193189 and SB431542 for 6 days. (**b**) Mesoderm differentiation was mediated by Activin A and Wnt3a treatment. TBX1 immunocytochemistry was performed after 2 days and GATA4 after 3 days. (**c**) Endoderm differentiation was also induced by Activin A and Wnt3a. Immunocytochemistry was performed after 3 days. hESC, H9 human embryonic stem cell line. Scale bars, 100 µm.

To confirm the finding that patient cells express a hypoglycosylated form of αDG, we carried out western blots on hiPSC culture lysates using the IIH6C4 antibody. Probing for glycosylated αDG showed a reduction that roughly correlated with the clinical severity for each patient (Fig. 3b, c). Control hiPSCs expressed αDG glycoforms averaging ∼140 kDa, which is slightly less than reported in human muscle [26]. Previous analysis suggests that low molecular weight forms of αDG – a consequence of fewer glycan structures – is a specific biochemical hallmark associated with disease severity [14]. Our study followed this pattern, with cells from the clinically mild FKRP patient expressing glycosylated αDG of the same mass as controls but in reduced abundance. Glycosylated αDG from the LARGE patient, who was of intermediate severity, showed a ∼30 kDa downward shift in molecular weight. hiPSCs from the severe POMT2 patient were virtually devoid of glycosylated αDG.

To test the functional impact of these hypoglycosylated forms of αDG, we performed a laminin overlay assay to measure the affinity of αDG for one of its ECM ligands, laminin. All patient cell lines showed reduced αDG-laminin binding activity that closely matched their degree of αDG hypoglycosylation (Fig. 3b, d). Blotting with antibodies against the core peptide of αDG (core-αDG) and β-dystroglycan (βDG) indicated similar expression across all samples, demonstrating that the dystroglycan proteins are expressed but that αDG is hypoglycosylated in dystroglycanopathy patient hiPSCs (Fig. 3b).

### Differentiation of Embryoid Bodies from Dystroglycanopathy hiPSCs

Muscle, eye, and brain abnormalities linked to basal lamina defects is a frequent finding in the dystroglycanopathies. Given that dystroglycanopathy patient hiPSCs exhibit the biochemical hallmark of the disease (i.e. hypoglycosylation of the basal lamina receptor αDG), we next asked whether patient embryoid bodies can synthesize basal lamina. To investigate potential disease-related phenotypes, we initially restricted our analysis to embryoid bodies from the LARGE patient and POMT2 patient, who had moderate and severe clinical findings respectively.

We differentiated control and patient hiPSCs into embryoid bodies using the protocol described earlier. Embryoid bodies from all cell lines contained a basal lamina sandwiched by epithelial and endodermal compartments (Fig. 5a). There was noticeable morphological variation across cell lines, possibly related to genetic background or clonal differences. Particularly in embryoid bodies from the third control and the LARGE patient, there was an occasional inversion of tissue layers such that the epithelial cells were on the exterior of the embryoid body (Fig. 5a).

**Fig. 5.**
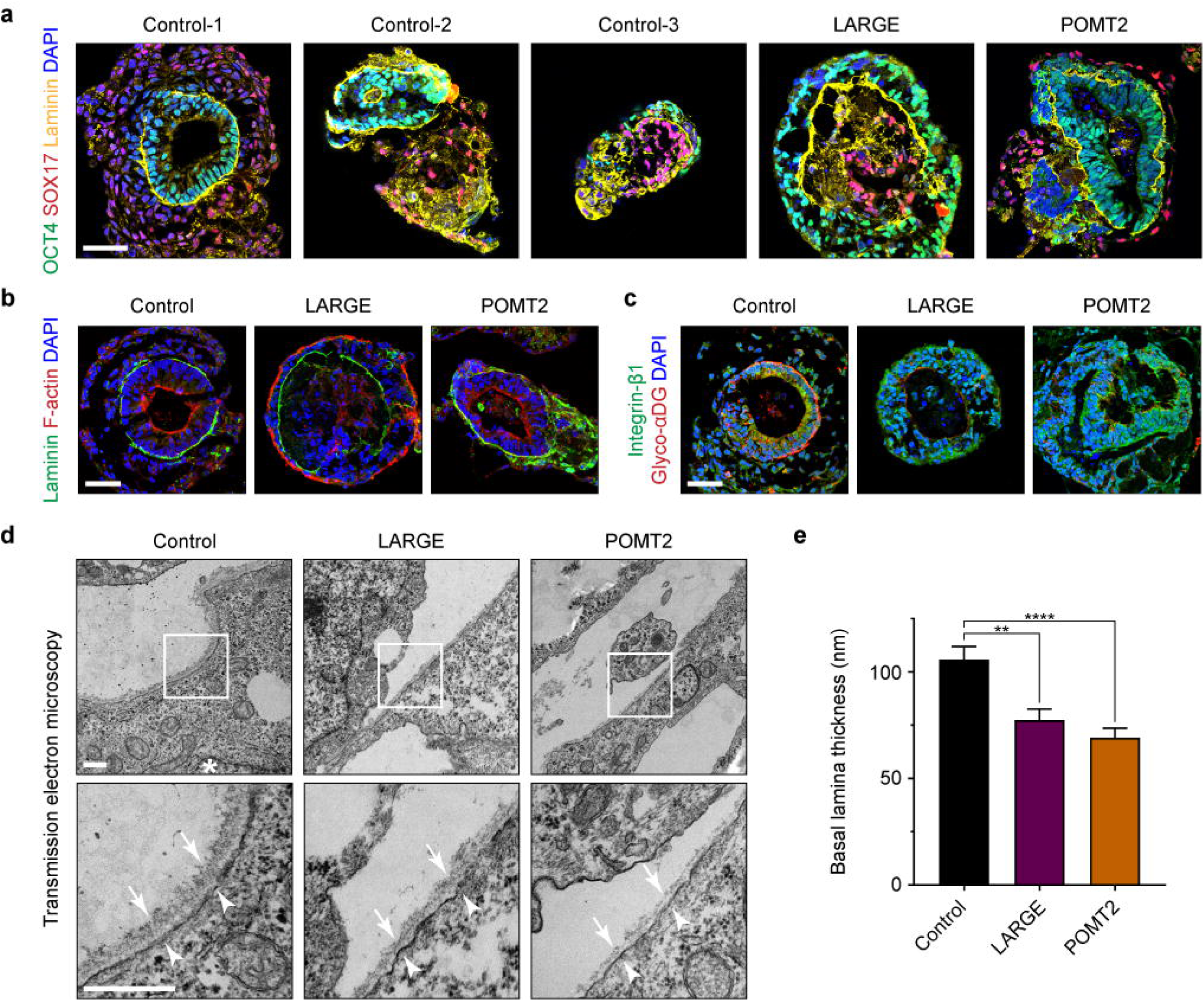
Ultrastructural Basal Lamina Defects in Dystroglycanopathy Patient Embryoid Bodies. (**a**-**c**) Representative immunohistochemistry images of control and patient embryoid bodies at day 5 of differentiation. At least three independent differentiations were carried out on each of three control hiPSC lines and two hiPSC clones from the LARGE and POMT2 patient. Scale bars, 50 µm. (**d**) Transmission electron micrographs of embryoid body basal lamina. Asterisk, nucleus; arrows, basal lamina; arrowheads, plasma membrane; scale bars, 500 µm. (**e**) Measurements of basal lamina thickness. Three control lines and two clones each from the LARGE and POMT2 patients were used. For each clone, micrographs of *n* ≥ 10 different basal lamina regions were collected across ≥ 5 embryoid bodies from 2-3 independent differentiations. Values displayed as mean ± s.e.m. Post-hoc comparisons ** P < 0.01, **** P < 0.0001.

Despite the heterogeneity between cultures, embryoid bodies from all control and patient hiPSC clones were similarly capable of assembling a laminin-rich basal lamina at the surface of the OCT4+ epithelium. Additionally, all embryoid bodies showed the expected morphology of epithelial polarity. OCT4+ cells were radially arranged and exhibited apicobasal polarity, with F-actin distributed on the cellular edge opposite from the basal lamina (Fig. 5b). Thus, at this resolution of analysis, we detected no major phenotypic difference between control and dystroglycanopathy embryoid bodies.

Consistent with our finding in undifferentiated hiPSCs, LARGE and POMT2 embryoid bodies were minimally reactive to an antibody recognizing glycosylated αDG. However, all embryoid bodies expressed the other major basal lamina receptor, integrin-β1 (Fig. 5c). Study of mouse embryoid bodies has shown functional redundancy between integrin-β1 and αDG in anchoring ECM molecules to the cell membrane [31]. This likely explains the ability of dystroglycanopathy patient embryoid bodies to assemble a basal lamina in the absence of αDG receptor function.

### Ultrastructural Basal Lamina Defects in Dystroglycanopathy Embryoid Bodies

Previous studies of dystroglycanopathy patient and mouse tissue have revealed ultrastructural ECM defects [14, 25]. To visualize embryoid body ECM ultrastructure, we employed transmission electron microscopy on thin sections of control and patient samples. In electron micrographs, basal lamina from control embryoid bodies was visible as a fibrous layer at the epithelial cell surface, roughly 100 nm thick (Fig. 5d). The overlying endodermal cell, some distance away, was always devoid of basal lamina. The basal lamina was composed of a ‘lamina lucida’ and a ‘lamina densa’ compartment. The lamina lucida – a thin electron-light layer at the epithelial plasma membrane – is believed to be spanned by the laminin long arm bound to its cell surface receptors, αDG and integrin [42]. The lamina densa – a thicker fibrous layer of electron-dense material just above the lamina lucida – is comprised of laminin cross-linking arms, perlecan, nidogen, and COLIV. However, the existence or extent of the lamina lucida may also be an artifact of our sample dehydration method [6].

The epithelial cells of embryoid bodies showed typical features of lateral, apical, and basal polarization. We observed electron-dense tight junctions at cell-cell borders, and microvilli decorated the epithelial cells’ apical aspect facing the embryoid body lumen (Fig. 6). Nuclei were polarized toward the basal aspect of epithelial cells in contact with the basal lamina (Fig. 5d). Occasionally, filamentous matrix could be seen in the extracellular space between endodermal and epithelial cells, but it was rarely attached to the basal lamina itself (Fig. 6). Therefore, the ECM structures in our embryoid bodies meet the criteria for an epithelial basal lamina. However, they cannot be categorized as a complete basement membrane, which requires an adjoined layer of ‘lamina reticularis’ fibrillar collagens [41].

**Fig. 6.**
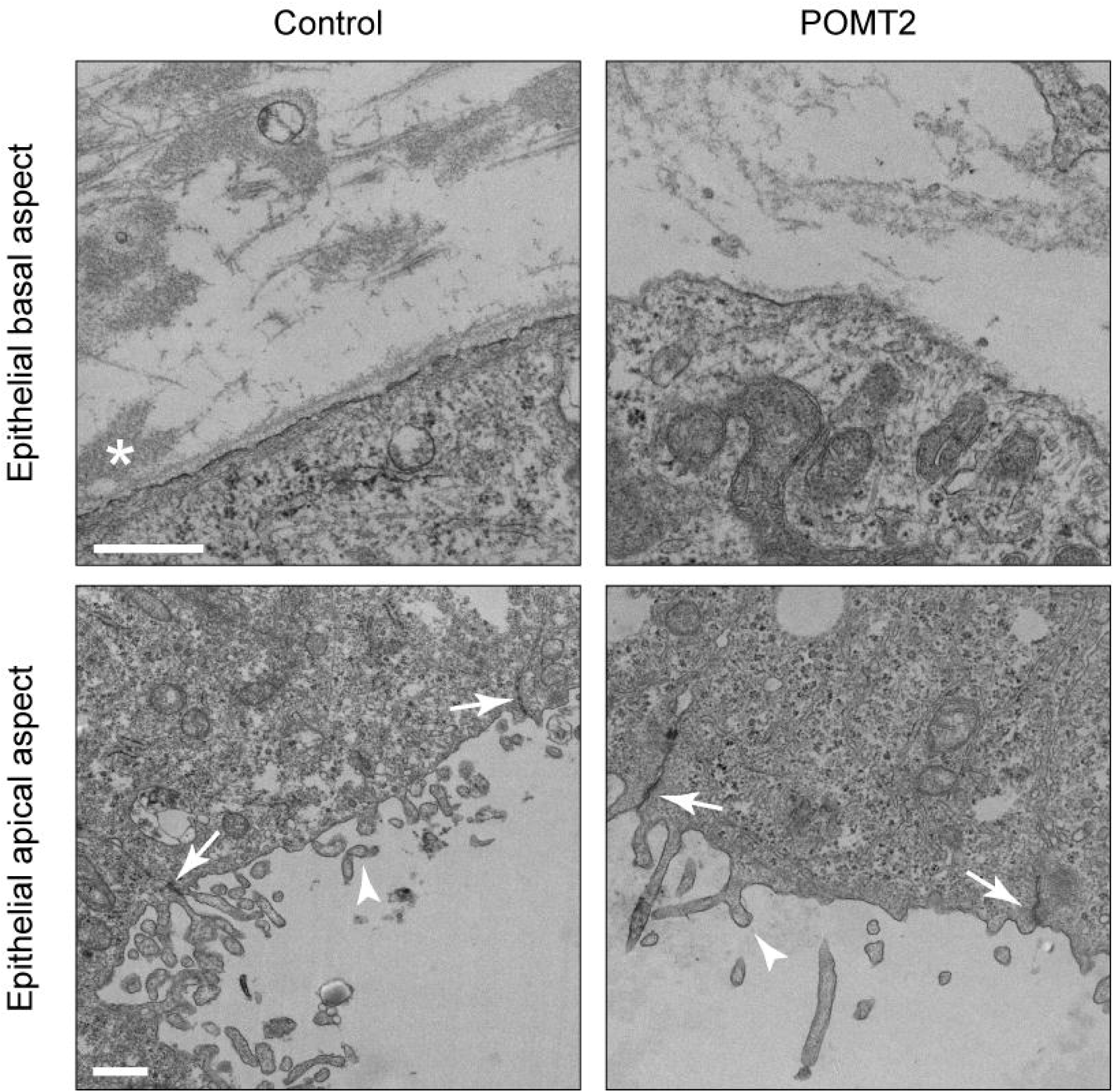
Ultrastructural Examination of Embryoid Body Morphology. Representative transmission electron micrographs depicting the ultrastructural morphology of embryoid body epithelium. Rare attachment of fibrillar matrix to the basal lamina indicated by asterisk. Arrows demonstrate epithelial tight junctions, and the arrow head indicates apical microvilli. Scale bar, 500 nm.

In LARGE and POMT2 embryoid bodies, the basal lamina was noticeably thinner due to a reduction of material in the lamina densa (Fig. 5d). Specifically, in POMT2 embryoid bodies, the basal lamina occasionally contained nanoscopic discontinuities (Fig. 5d). The basal lamina in control embryoid bodies measured 105.8 ± 6.1 nm thick. LARGE and POMT2 basal lamina were significantly thinner at 77.5 ± 5.1 and 69.1 ± 4.5 nm respectively (mean ± s.e.m., P = 0.0022 and P < 0.0001) (Fig. 5e).

We considered whether a deficit of certain ECM molecules might explain the reduced thickness of patient basal lamina. In addition to laminin, αDG directly binds to perlecan, which in turn crosslinks nidogen and collagen type IV (COLIV) to form the basal lamina [39, 44]. Overall, there were no consistently detectable differences in the staining pattern or intensity of these basal lamina components between control and patient embryoid bodies (Fig. 7a). However, in some POMT2 embryoid bodies, there was occasionally a subtle loss of perlecan and COLIV co-localization with laminin (Fig. 7b).

**Fig. 7.**
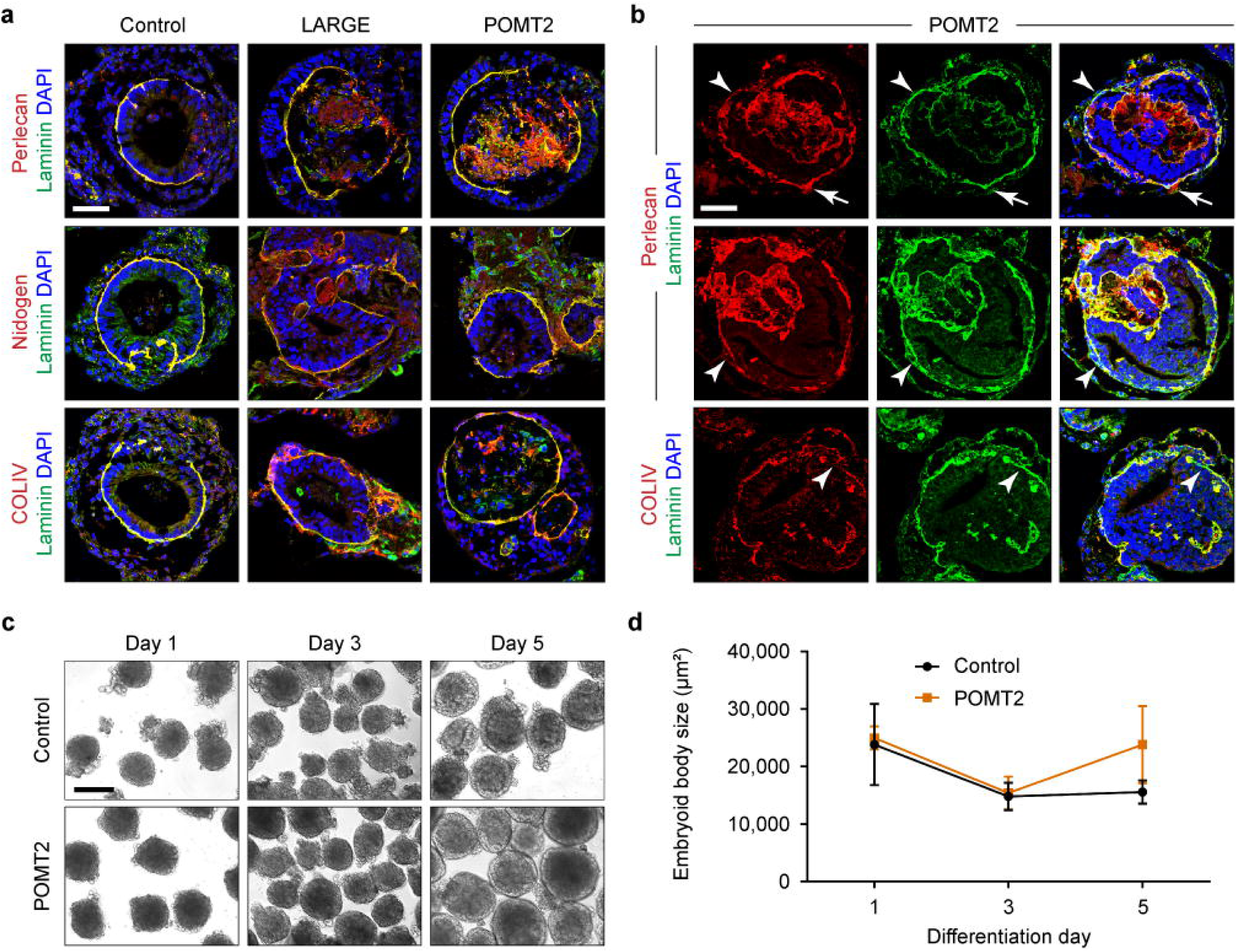
Analysis of Basal Lamina Components and Growth Characteristics of Dystroglycanopathy Embryoid Bodies. (**a**) Representative antibody labeling to assess co-localization of laminin with other basal lamina constituents. Embryoid bodies from three controls and two clones per patient were used. Scale bar, 50 µm. (**b**) Examples of occasionally separate localization of laminin, perlecan, and COLIV in POMT2 embryoid bodies. The arrow shows localization of perlecan outside the basal lamina. Arrowheads indicate the presence of laminin without perlecan or COLIV. Scale bar, 50 µm. (**c**) Phase-contrast images of embryoid body differentiation on days 1, 3, and 5. Scale bar, 100 µm. (**d**) Quantification of embryoid body size over time based on averaged cross-sectional area measurements in phase-contrast images from *n* = 3 differentiations. At least 75 embryoid bodies per line were analyzed during each differentiation. No statistical significance was found between control and patient at any time point (P > 0.05), values expressed as mean ± s.d.

Over the course of differentiation, embryoid bodies were remarkably static in size, and there was no significant difference between control and POMT2 embryoid body surface area (Fig. 7c, d). This contrasts greatly with brain and muscle tissue, which undergo significant size expansion and mechanical strain during embryonic development and muscle contraction, respectively [20, 24]. Thus, our embryoid body differentiation protocol serves as a simplified model for patient-specific basal lamina assembly, without the additional variables of tissue movement and growth. In this system, dystroglycanopathy embryoid bodies with mutations in *LARGE* or *POMT2* can synthesize a basal lamina of apparently typical molecular composition but with abnormal ultrastructure. These data corroborate published results demonstrating that αDG is generally dispensable for initial basal lamina formation [9, 30, 35].

Because our embryoid body differentiation protocol requires co-culture of hiPSCs with MEFs, we reasoned that abundant fibroblast ECM secretion could be masking disease-related deficits in basal lamina assembly. To evaluate this possibility, we labeled control and POMT2 hiPSCs and embryoid bodies with antibodies against human nuclear antigen (HuNu). In feeder-dependent hiPSC cultures, human hiPSCs were clearly distinguished by HuNu expression, while MEFs were heavily labeled by laminin antibodies (Fig. 8a). In embryoid bodies, hiPSCs and MEFs self-segregated within 24 hours of spheroid formation, with MEFs budding off and ultimately detaching from the differentiating embryoid body 2 – 4 days before basal lamina formation (Fig. 8b). Because all analyses were conducted on day 5 embryoid bodies, we believe the presence of MEFs is not a significant confounding factor in our experiments.

**Fig. 8.**
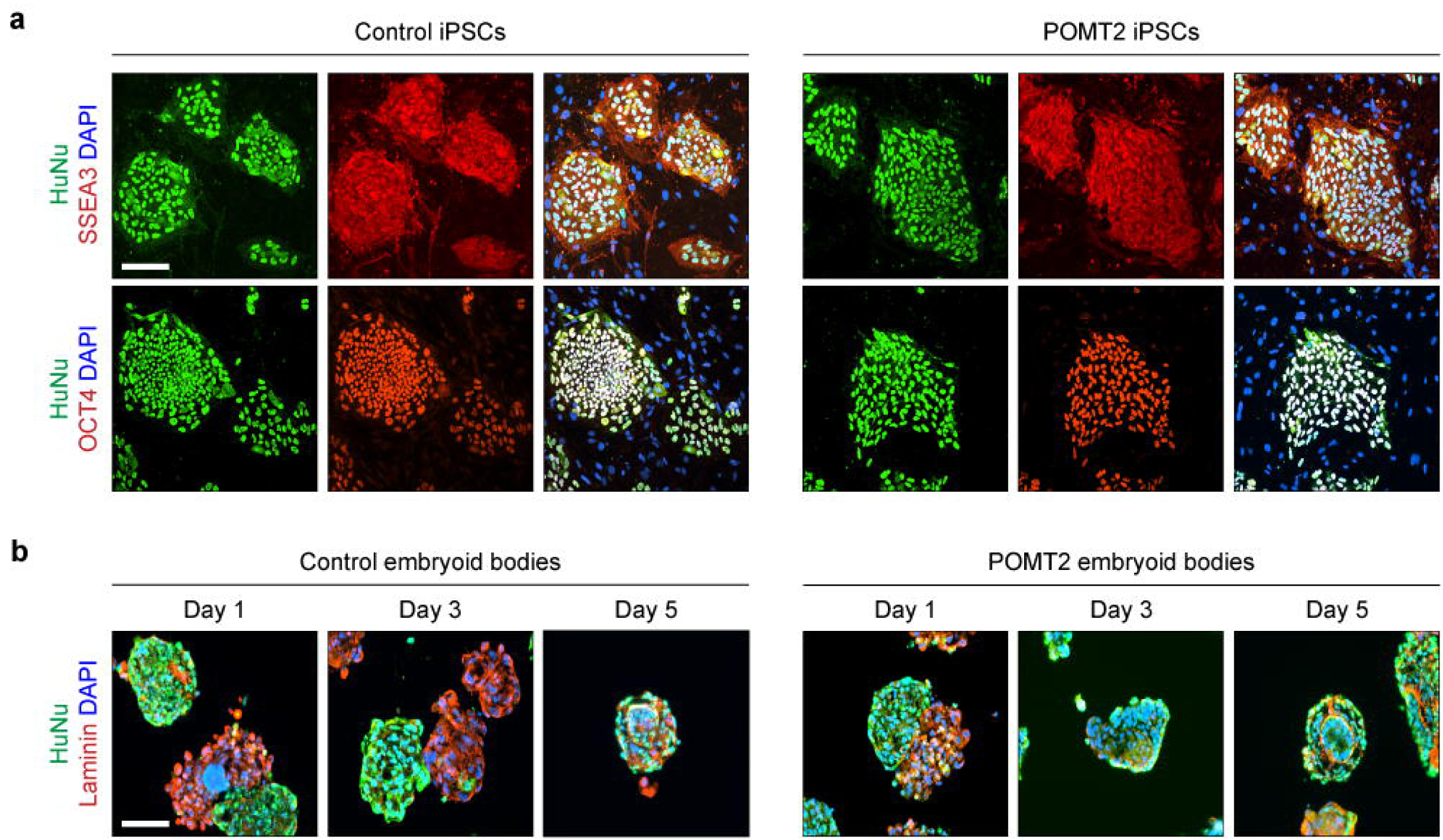
Morphologically Mature Embryoid Bodies are Virtually Devoid of MEFs. (**a**) Immunocytochemistry of feeder-dependent hiPSCs to distinguish human cells (HuNu) from MEFs. Control-1 hiPSCs and one hiPSC clone of the POMT2 patient were used. (**b**) Embryoid bodies at different time points, derived from control and POMT1 hiPSCs. Scale bars, 100 µm.

### Impaired Basal Lamina Assembly on Endoderm-Free Dystroglycanopathy Embryoid Bodies

Knockout of dystroglycan in mouse embryonic stem cells and neural stem cells has been reported to reduce laminin polymerization at the cell surface, which is a prerequisite for basal lamina formation [21, 52]. Because we observed thinner basal lamina in dystroglycanopathy patient embryoid bodies, we next investigated whether this phenotype might be linked to a reduced ability to polymerize laminin. We developed an embryoid body culture protocol that both prevents the formation of laminin-secreting endoderm and obviates the need for MEF co-culture (Fig. 9a, b). This eliminated the major sources of ECM and allowed for precise control of laminin concentration by exogenous supplementation in the growth medium.

**Fig. 9.**
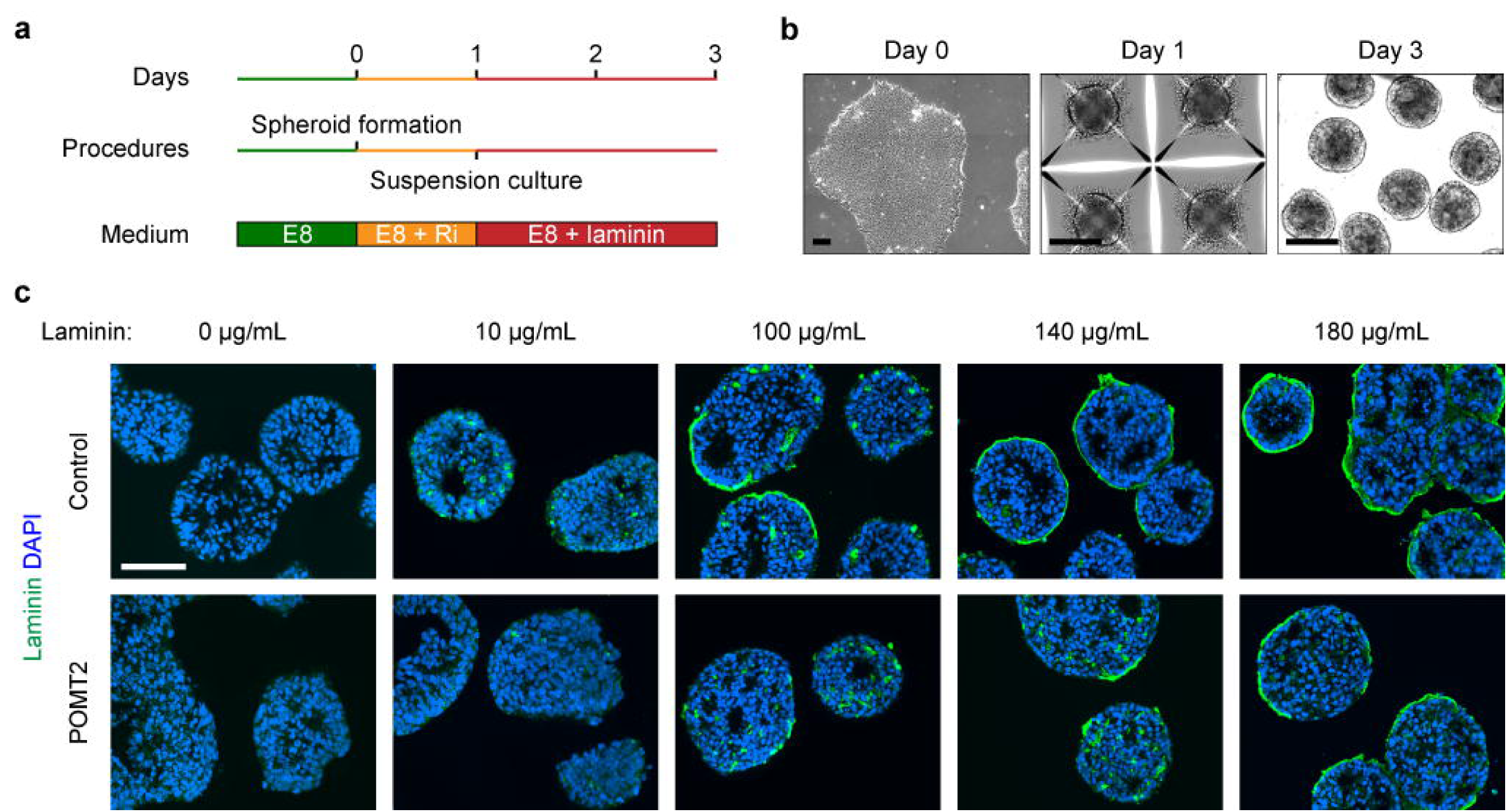
Assembly of Exogenous Laminin on Endoderm-Free Embryoid Bodies. (**a**, **b**) Schematic and phase-contrast of endoderm-free embryoid body culture protocol. Feeder-free hiPSCs were seeded on day 0 in microwell plates to form spheroids of roughly 1,000 cells by day 1. The spheroids were transferred to suspension culture supplemented with laminin for 48 hours. (**c**) Immunohistochemistry demonstrating the effect of increasing laminin concentration on endoderm-free embryoid bodies. Embryoid bodies were supplemented with varying concentrations of laminin on day 1 and collected for analysis 48 hours later. Scale bars, 100 µm.

In the absence of laminin, endoderm-free embryoid bodies self-assembled into disorganized, cavitated spheroids (Fig. 9c). We supplemented various concentrations of laminin to control embryoid bodies and inspected the cultures after 48 hours. Amounts greater than 100 µg/mL resulted in a thin layer of laminin at the surface of some embryoid bodies. In embryoid bodies with significant laminin recruitment, the central cavity was widened, and cells with direct laminin contact adopted a polarized orientation (Fig. 9c). POMT2 embryoid bodies – from the most severe dystroglycanopathy patient – showed a striking absence of accumulated surface laminin except in the highest concentration tested, 180 µg/mL. To avoid ceiling and floor effects, we used 140 µg/mL laminin in subsequent experiments, because approximately half of the control embryoid body surface area bound laminin at this concentration (Fig. 9c).

We tested endoderm-free embryoid bodies from control subjects and LARGE, FKRP, and POMT2 dystroglycanopathy patients for their capacity to assemble laminin (Fig. 10a). Compared to controls, which accumulated laminin on 65.7 ± 6.3% of surfaces, POMT2 embryoid bodies showed 20.5 ± 5.2% surface laminin (mean ± s.e.m., P = 0.0006) (Fig. 10d). In contrast, LARGE and FKRP embryoid bodies assembled 48.4 ± 7.3% and 46.1 ± 10% laminin respectively, which was also reduced but not statistically different from controls (P = 0.405 and P = 0.283) (Fig. 10d).

**Fig. 10.**
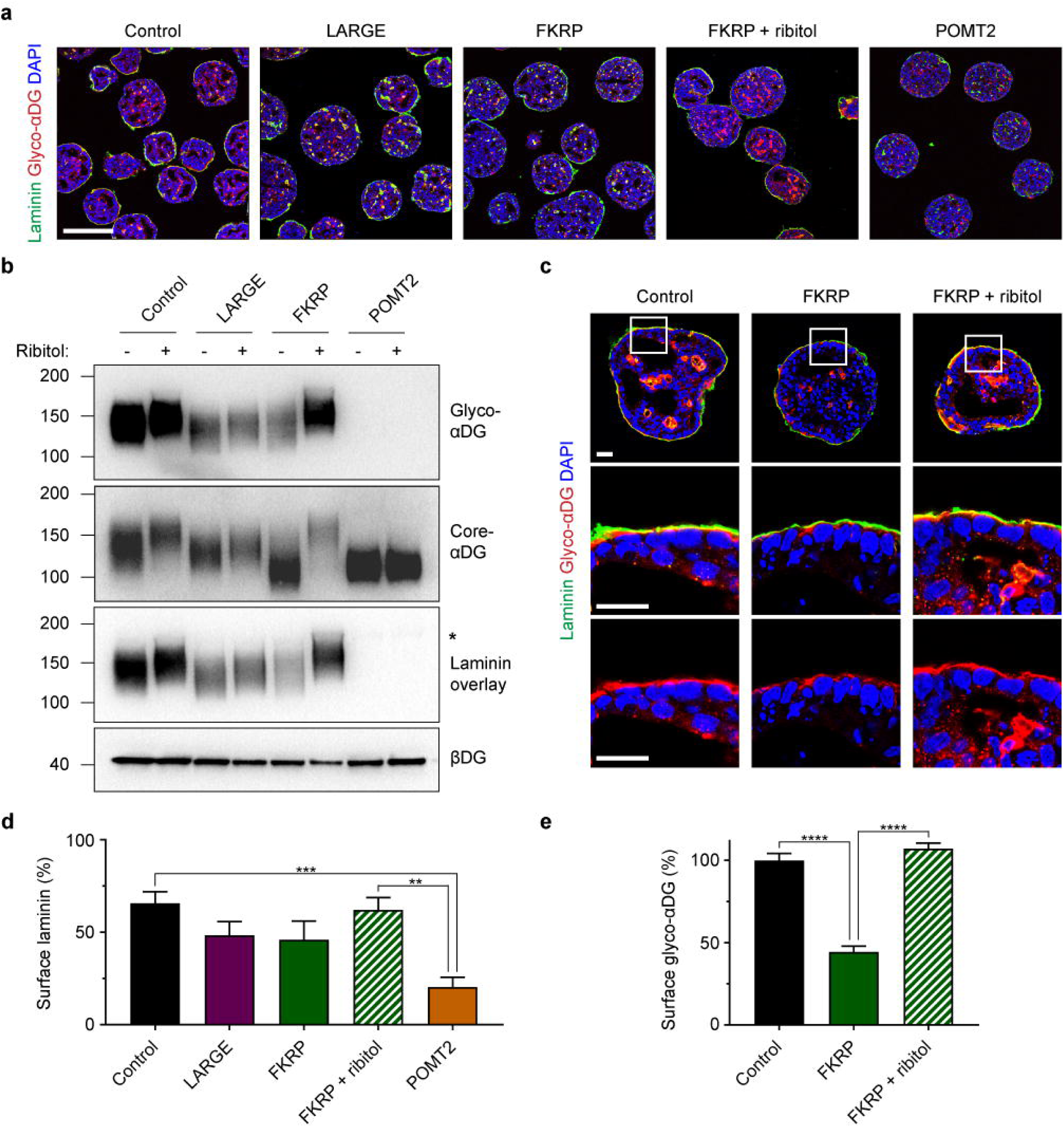
Patient-Specific Differences in Laminin Assembly and Response to Ribitol Treatment. (**a**) Immunohistochemistry on day 3 endoderm-free embryoid bodies. Scale bar, 200 µm. (**b**) Western blotting of protein lysates from hiPSC cultures. hiPSCs were supplemented with daily medium changes with (+) or without (-) 3 mM ribitol for 72 hours before protein was collected. Asterisk indicates the position of endogenous laminin. (**c**) Immunohistochemistry on day 3 endoderm-free embryoid bodies to assess the effect of ribitol treatment on the localization and glycosylation of αDG in the FKRP patient. (**d**) Quantification of the percent embryoid body surface area covered by laminin. Three controls and two clones per patient were analyzed across *n* = 3 differentiations each. Values plotted as mean ± s.e.m. Post-hoc comparisons ** P < 0.01, *** P < 0.001. (**e**) Quantification of glycosylated αDG staining intensity at the embryoid body surface. Three controls and two FKRP clones with and without ribitol were analyzed and normalized to a percentage of the controls. For each clone, *n* = 27 surface regions of 50 µm length were quantified across three differentiations. Values graphed as mean ± s.e.m. Post-hoc comparisons **** P < 0.0001.

### Ribitol Treatment Promotes Functional Glycosylation of αDG in FKRP Patient Embryoid Bodies

FKRP is a glycosyltransferase enzyme that catalyzes the addition of ribitol phosphate to αDG [13, 27, 40]. This is an essential enzymatic step for the post-translational installation of matriglycans on αDG, which are the structural basis for αDG-ligand binding [4]. Recently, dietary supplementation with ribitol was shown to improve muscle phenotypes in a mouse model of FKRP-related dystroglycanopathy [5].

To determine the effect of ribitol on dystroglycanopathy patient hiPSCs, we supplemented the culture medium with 3 mM ribitol daily and harvested the cells 72 hours later. This dosage was based on previous reports of efficacious treatment with 3mM ribitol or CDP-ribitol in ISPD mutant dystroglycanopathy cell cultures [13, 27]. Ribitol treatment of FKRP hiPSCs greatly increased the abundance of glycosylated αDG and resulted in a ∼20 kDa upward shift in molecular weight (Fig. 10b, 11a). This change was accompanied by a comparable increase in αDG-laminin binding. Ribitol-treated LARGE hiPSCs were unchanged in glycosylated αDG quantity but showed a slight increase in molecular weight (5-10 kDa). As would be expected, POMT2 hiPSCs, in which glycosylated αDG is essentially absent, showed no improvement with ribitol exposure (Fig. 10b). In addition to ribitol phosphate, FKRP is known to transfer glycerol phosphate onto αDG, which inhibits its functional glycosylation. However, we found that treating control and FKRP hiPSCs with 3 mM glycerol daily for 72 hours had no discernable effect on αDG glycosylation (Fig. 11b).

**Fig. 11.**
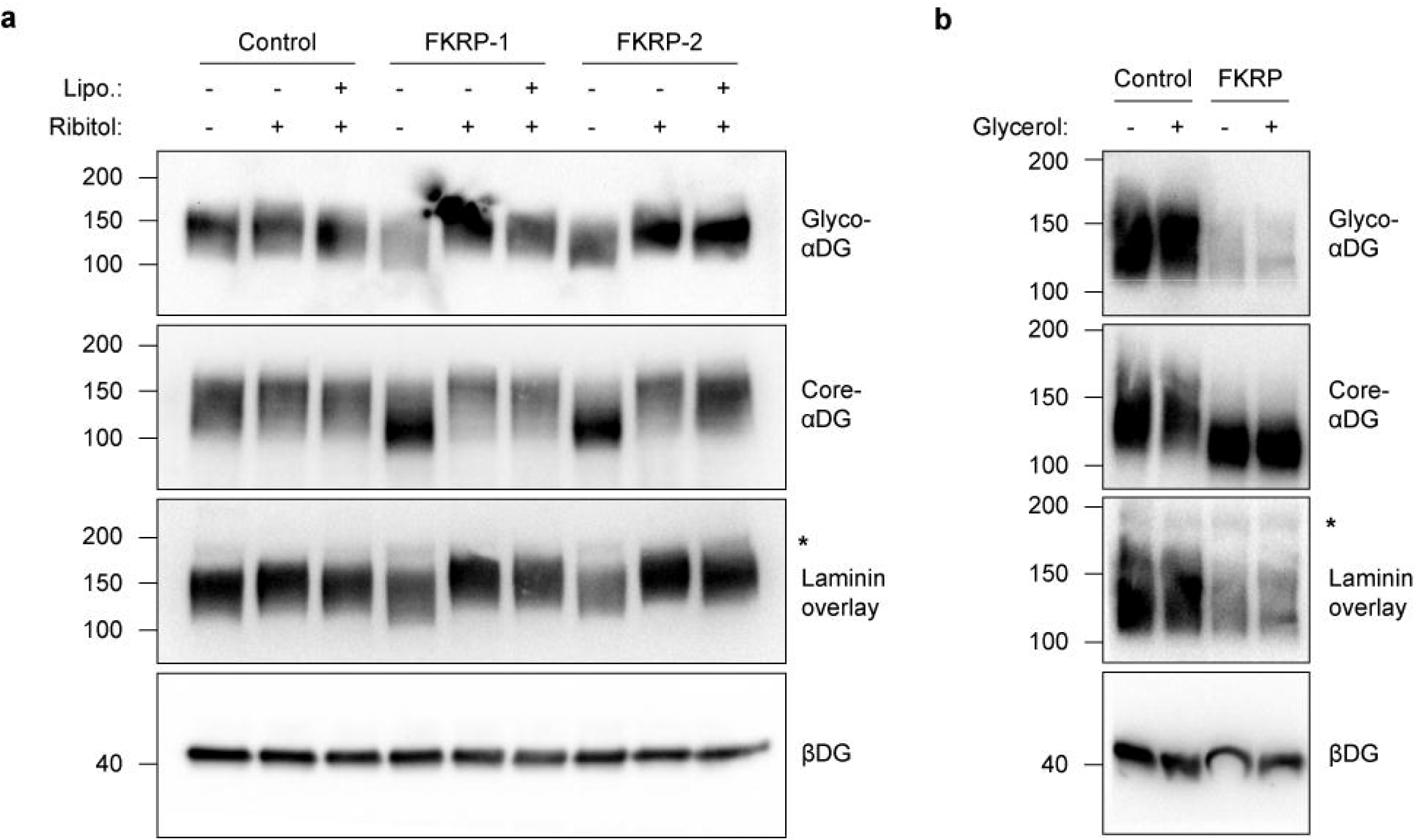
Ribitol Treatment Promotes Functional Glycosylation of αDG in FKRP hiPSCs. (**a**) Additional western blots on control and FKRP patient hiPSCs. Cells were supplemented with 3 mM ribitol in daily medium changes for 72 hours before protein was harvested. Asterisks indicate position of endogenous laminin in the samples. Lipofectamine (Lipo.) (STEM00003, Thermo Fisher) was tested in conjunction with ribitol treatment to enhance delivery to cells. In this condition, ribitol and lipofectamine were administered for only 24 hours following manufacturer’s instructions, and the cells were collected 72 hours later. No apparent difference was observed between 24-hour lipofectamine-delivered and 72 hour free-uptake of ribitol. (**b**) Daily administration of 3 mM glycerol for 72 hours shows no effect on αDG by western blot.

Because ribitol treatment resulted in a specific and profound improvement to glycosylated αDG in FKRP hiPSCs, its effect was next assessed in FKRP endoderm-free embryoid bodies. hiPSCs were treated with 3 mM ribitol daily starting 72 hours before embryoid body formation and continued throughout the experiment. Treatment of FKRP embryoid bodies lead to a noticeable increase in the staining intensity of glycosylated αDG (Fig. 10a, c). There was a slight but non-significant increase in surface laminin polymerization on ribitol treated FKRP embryoid bodies (62.1 ± 6.6%) compared to untreated FKRP embryoid bodies (46.1 ± 10%; mean ± s.e.m, P = 0.566) (Fig. 10d). Importantly though, at laminin contact points, glycosylated αDG was upregulated to the same level as controls (ribitol-treated FKRP: 107.2 ± 3.4%; untreated FKRP: 44.5 ± 3.5%; normalized to control glyco-αDG, mean ± s.e.m., P < 0.0001) (Fig. 10e). These results demonstrate ribitol treatment as a viable therapeutic strategy for upregulating αDG glycosylation in FKRP patient tissue.

## Discussion

Our results provide an approach for evaluating patient-specific basal lamina *in vitro*. We applied this method to study patients with dystroglycanopathy. We found that patient hiPSC-derived embryoid bodies recapitulate the clinical disease spectrum and exhibit ECM defects seen in animal models. Finally, this system allowed us to evaluate the efficacy of ribitol supplementation – a recent candidate therapeutic for the dystroglycanopathies – in patient samples of different genotypes.

Embryoid body-based methods, which essentially involve culturing pluripotent stem cells in 3D aggregates, are widely used in stem cell research. Typical applications include evaluating the pluripotency of new cell lines or as an intermediate stage during differentiation toward specific lineages [28]. Under strict growth conditions, embryoid bodies from mouse and human embryonic stem cells (ESCs) can exquisitely self-organize into structures mimicking the pre-gastrulation embryo. These have served as models for early embryonic events – epiblast polarization and ECM formation – that are required for tissue genesis [29].

Our human hiPSC-derived embryoid bodies resemble their ESC-derived counterparts by similarly requiring serum-containing media and MEF co-culture to form polarized epiblast, basal lamina, and extra-embryonic endoderm. It remains unclear why these conditions uniquely enable such self-organization. BMPs, which are abundant in serum [22], are known to induce extra-embryonic endoderm differentiation [16]. Co-culture with MEFs can also influence the lineage bias of ESCs [32]. Thus, it may be a combination of soluble factors from serum and MEFs that promotes the simultaneous existence of endoderm and epiblast cell populations in our hiPSC-derived embryoid bodies.

Dual genetic deletion of αDG and integrin-β1 prevents basement membrane formation and epithelial polarization in mouse ESC-derived embryoid bodies. Because these two receptors have overlapping roles, expression of one can partly rescue the function of the other [31]. By studying dystroglycanopathy patient embryoid bodies, we extend these findings to show that a moderate reduction in αDG receptor activity – with normal expression of both αDG and integrin-β1 core proteins – is sufficient to cause subtle ECM deficits that may underlie disease pathogenesis.

A thin and discontinuous basal lamina has been reported in the muscle of dystroglycanopathy patients [25, 48], similar to our observation in patient embryoid bodies. Also, the retina inner limiting basement membrane is thin, patchy, and less stiff in *Pomgnt1*-null mice [23, 52]. One caveat of our hiPSC-derived embryoid body system is that it seems to only recapitulate basal lamina development. It is unclear how the observed basal lamina defects might translate to the formation of a full basement membrane. Furthermore, the specific structural and molecular deficits underlying such abnormally thin basal lamina are still unknown.

In contrast to the above findings, basement membranes in dystroglycan-null mouse embryoid bodies, and in the muscle of *Large* mutant mice and certain human patients, are thicker than controls, sometimes with mislocalized ECM components [14, 30]. These dichotomous observations may be related to differences in ECM recruitment, organization, and maintenance at the cell surface, as a consequence of reduced αDG receptor activity but under specific tissue and disease state circumstances. One study reported a mixture of thinned and duplicated basal lamina in different regions of dystroglycanopathy patient muscle [46]. Therefore, whether the diseased basal lamina is thin and patchy, or thick and duplicated, may vary due to local tissue conditions.

The pathogenic mechanism of brain malformation in the dystroglycanopathies is still not fully understood. During brain development, the human cerebral cortex undergoes massive expansion and folding in the third trimester [12]. This rapid increase of surface area likely places mechanical strain on the brain’s basement membrane casing, necessitating timely ECM remodeling to accommodate the larger area. In mice, genetic deletion of ECM genes or hypoglycosylation of αDG both result in basement membrane rupture and ensuing tissue malformation during this most rapid phase of brain development [2, 19, 37, 43]. These data present αDG as one of several laminin receptors that contributes to the efficient organization of ECM molecules into a coherent basement membrane.

Our result – that endoderm-free patient embryoid bodies show diminished laminin accumulation – corroborates evidence that αDG-deficient cells have reduced ECM-binding kinetics that might render basement membranes susceptible to mechanically-induced deformation [21, 52]. Crucially, we found that embryoid bodies undergo minimal volumetric growth. This could explain the relatively mild phenotypes in embryoid bodies of even the most severe dystroglycanopathy patient.

The human brain specifically undergoes greatest expansion in the occipital, temporal, and lateral parietal cortices [12]. Such regional growth dynamics suggest one possible basis for the spatial arrangement of the typical cortical malformation in the dystroglycanopathies, which are fundamentally due to breaches in basement membrane integrity, resulting in cellular over-migration beyond the confines of the [1, 7, 33, 50]. These particular growth dynamics may also underlie some of the differences between human patients and mouse models [3].

There is currently no effective treatment for the dystroglycanopathies, and the diversity of underlying genetic causes for the disease presents a challenge for targeted therapies. Supplementation of the sugar alcohol ribitol has recently emerged as a promising therapeutic for specific classes of dystroglycanopathy. In mammalian cells, the enzyme ISPD synthesizes CDP-ribitol from ribitol, which is then attached to αDG by the glycosyltransferases FKTN and FKRP [13, 27, 40]. This enzymatic process is a critical step in constructing the laminin-binding glycan of αDG.

Treatment of ISPD mutant cells with ribitol or CDP-ribitol promotes αDG glycosylation [13, 27], and dietary ribitol supplementation rescues muscle phenotypes in Fkrp mutant mice [5]. Here, we extend this concept from animal models to a human patient-derived system. We found specific efficacy of ribitol on FKRP patient hiPSCs, as evidenced by a complete rescue of glycosylated laminin-binding αDG. There was minimal effect on LARGE and no effect on POMT2 patient hiPSCs, as would be expected from the location of these genetic forms in the pathway of αDG glycosylation. In FKRP endoderm-free embryoid bodies, ribitol significantly upregulated glycosylated αDG at the laminin interface. There was also slightly increased accumulation of laminin at the embryoid body surface, but this *in vitro* system may lack the complexity or sensitivity to further detect functional improvements in an already relatively mild patient.

The FKRP patient in this study harbors a L276I mutation, the most common variant in the FKRP-related dystroglycanopathies [11]. We speculate that ribitol supplementation may be a rational and effective treatment, in particular for mild-to-moderate FKRP and FKTN patients with residual ribitol transferase function that can be boosted by the additional supply of substrate. It remains to be investigated whether ribitol would also benefit additional groups of dystroglycanopathy patients. Collectively, these data establish a system to interrogate basal lamina structure and ECM receptor function in patient tissue, expanding the options for personalized phenotyping and drug evaluation in the dystroglycanopathies.

## Acknowledgements

We would like to thank the patients and their families for their participation. We are grateful to Susan Cheng, Virginia Crocker, and Sandra Lara for their EM technical support. Electron microscopy was performed in the NINDS EM facility. This work was funded by the US National Institutes of Health Intramural Research Program in the National Institute of Neurological Disorders and Stroke.

## Author contributions

ARN, KZ, and CGB conceived and designed the study. ARN performed all experiments. MML contributed to data collection and image analyses. BSM assisted in developing the embryoid body protocol. ARN wrote the manuscript and all authors edited the manuscript.

## References

1. Aida N, Tamagawa K, Takada K, Yagishita A, Kobayashi N, Chikumaru K, Iwamoto H (1996) Brain MR in Fukuyama congenital muscular dystrophy. Am J Neuroradiol 17:605–13

2. Booler HS, Pagalday-Vergara V, Williams JL, Hopkinson M, Brown SC (2016) Evidence of early defects in Cajal Retzius cell localisation during brain development in a mouse model of dystroglycanopathy. Neuropathol Appl Neurobiol 43:330–345. doi: 10.1111/ijlh.12426

3. Booler HS, Williams JL, Hopkinson M, Brown SC (2015) Degree of Cajal-Retzius cell mislocalisation correlates with the severity of structural brain defects in mouse models of dystroglycanopathy. Brain Pathol 26:465–478. doi: 10.1111/bpa.12306

4. Briggs D., Strazzulli A, Moracci M, Yu L, Yoshida-Moriguchi T, Zheng T, Venzke D, Anderson M, Hohenester E, Campbell KP (2016) Structural basis of laminin binding to the LARGE glycans on dystroglycan. Nat Chem Biol 12:810–814. doi: 10.2210/PDB5IK7/PDB

5. Cataldi MP, Lu P, Blaeser A, Lu QL (2018) Ribitol restores functionally glycosylated α-dystroglycan and improves muscle function in dystrophic FKRP-mutant mice. Nat Commun 9:3448. doi: 10.1038/s41467-018-05990-z

6. Chan FL, Inoue S (1994) Lamina lucida of basement membrane: An artefact. Microsc Res Tech 28:48–59. doi: 10.1002/jemt.1070280106

7. Clement E, Mercuri E, Godfrey C, Smith J, Robb S, Kinali M, Straub V, Bushby K, Manzur A, Talim B, Cowan F, Quinlivan R, Klein A, Longman C, McWilliam R, Topaloglu H, Mein R, Abbs S, North K, Barkovich AJ, Rutherford M, Muntoni F (2008) Brain involvement in muscular dystrophies with defective dystroglycan glycosylation. Ann Neurol 64:573–582. doi: 10.1002/ana.21482

8. Clement EM, Feng L, Mein R, Sewry CA, Robb SA, Manzur AY, Mercuri E, Godfrey C, Cullup T, Abbs S, Muntoni F (2012) Relative frequency of congenital muscular dystrophy subtypes: Analysis of the UK diagnostic service 2001-2008. Neuromuscul Disord 22:522–527. doi: 10.1016/j.nmd.2012.01.010

9. Cote PD, Moukhles H, Lindenbaum M, Carbonetto S (1999) Chimaeric mice deficient in dystroglycans develop muscular dystrophy and have disrupted myoneural synapses. Nat Genet 23:338–342. doi: 10.1038/15519

10. Devisme L, Bouchet C, Gonzalès M, Alanio E, Bazin A, Bessières B, Bigi N, Blanchet P, Bonneau D, Bonnières M, Bucourt M, Carles D, Clarisse B, Delahaye S, Fallet-Bianco C, Figarella-Branger D, Gaillard D, Gasser B, Delezoide A-L, Guimiot F, Joubert M, Laurent N, Laquerrière A, Liprandi A, Loget P, Marcorelles P, Martinovic J, Menez F, Patrier S, Pelluard F, Perez M-J, Rouleau C, Triau S, Attié-Bitach T, Vuillaumier-Barrot S, Seta N, Encha-Razavi F (2012) Cobblestone lissencephaly: neuropathological subtypes and correlations with genes of dystroglycanopathies. Brain 135:469–482. doi: 10.1093/brain/awr357

11. Frosk P, Greenberg CR, Tennese AAP, Lamont R, Nylen E, Hirst C, Frappier D, Roslin NM, Zaik M, Bushby K, Straub V, Zatz M, De Paula F, Morgan K, Fujiwara TM, Wrogemann K (2005) The most common mutation in FKRP causing limb girdle muscular dystrophy type 21 (LGMD2I) may have occurred only once and is present in Hutterites and other populations. Hum Mutat 25:38–44. doi: 10.1002/humu.20110

12. Garcia KE, Robinson EC, Alexopoulos D, Dierker DL, Glasser MF, Coalson TS, Ortinau CM, Rueckert D, Taber LA, Van Essen DC, Rogers CE, Smyser CD, Bayly P V. (2018) Dynamic patterns of cortical expansion during folding of the preterm human brain. Proc Natl Acad Sci 115:3156–3161. doi: 10.1073/pnas.1715451115

13. Gerin I, Ury B, Breloy I, Bouchet-Seraphin C, Bolsée J, Halbout M, Graff J, Vertommen D, Muccioli GG, Seta N, Cuisset J-M, Dabaj I, Quijano-Roy S, Grahn A, Van Schaftingen E, Bommer GT (2016) ISPD produces CDP-ribitol used by FKTN and FKRP to transfer ribitol phosphate onto α-dystroglycan. Nat Commun 7:11534. doi: 10.1038/ncomms11534

14. Goddeeris MM, Wu B, Venzke D, Yoshida-Moriguchi T, Saito F, Matsumura K, Moore SA, Campbell KP (2013) LARGE glycans on dystroglycan function as a tunable matrix scaffold to prevent dystrophy. Nature 503:136–140. doi: 10.1038/nature12605

15. Godfrey C, Foley AR, Clement E, Muntoni F (2011) Dystroglycanopathies: coming into focus. Curr Opin Genet Dev 21:278–285. doi: 10.1016/j.gde.2011.02.001

16. Graham SJL, Wicher KB, Jedrusik A, Guo G, Herath W, Robson P, Zernicka-Goetz M (2014) BMP signalling regulates the pre-implantation development of extra-embryonic cell lineages in the mouse embryo. Nat Commun 5:5667. doi: 10.1038/ncomms6667

17. Graziano A, Messina S, Bruno C, Pegoraro E, Magri F (2015) Prevalence of congenital muscular dystrophy in Italy: A population study. Neurology 84:904–911. doi: 10.1212/WNL.0000000000001303

18. Grunseich C, Zukosky K, Kats IR, Ghosh L, Harmison GG, Bott LC, Rinaldi C, Chen K, Chen G, Boehm M, Fischbeck KH (2014) Stem cell-derived motor neurons from spinal and bulbar muscular atrophy patients. Neurobiol Dis 70:12–20. doi: 10.1016/j.nbd.2014.05.038

19. Halfter W, Dong S, Yip Y-P, Willem M, Mayer U (2002) A critical function of the pial basement membrane in cortical histogenesis. J Neurosci 22:6029–6040. doi: 20026580

20. Han R, Kanagawa M, Yoshida-Moriguchi T, Rader EP, Ng RA, Michele DE, Muirhead DE, Kunz S, Moore SA, Iannaccone ST, Miyake K, McNeil PL, Mayer U, Oldstone MBA, Faulkner JA, Campbell KP (2009) Basal lamina strengthens cell membrane integrity via the laminin G domain-binding motif of α-dystroglycan. Proc Natl Acad Sci 106:12573–12579. doi: 10.1073/pnas.0906545106

21. Henry MD, Campbell KP (1998) A Role for Dystroglycan in Basement Membrane Assembly. Cell 95:859–870. doi: 10.1016/S0092-8674(00)81708-0

22. Herrera B, Inman GJ (2009) A rapid and sensitive bioassay for the simultaneous measurement of multiple bone morphogenetic proteins. Identification and quantification of BMP4, BMP6 and BMP9 in bovine and human serum. BMC Cell Biol 10:20. doi: 10.1186/1471-2121-10-20

23. Hu H, Candiello J, Zhang P, Ball SL, Cameron D a, Halfter W (2010) Retinal ectopias and mechanically weakened basement membrane in a mouse model of muscle-eye-brain (MEB) disease congenital muscular dystrophy. Mol Vis 16:1415–1428

24. Hu H, Yang Y, Eade A, Xiong Y, Qi Y (2007) Breaches of the Pial Basement Membrane and Disappearance of the Glia Limitans During Development Underlie the Cortical Lamination Defect in the Mouse Model of Muscle-Eye-Brain Disease. J Comp Neurol 502:168–183. doi: 10.1002/cne

25. Ishii H, Hayashi YK, Nonaka I, Arahata K (1997) Electron microscopic examination of basal lamina in Fukuyama congenital muscular dystrophy. Neuromuscul Disord 7:191–7

26. Jensen BS, Willer T, Saade DN, Cox MO, Mozaffar T, Scavina M, Stefans VA, Winder TL, Campbell KP, Steven A, Mathews KD (2015) GMPPB-Associated Dystroglycanopathy: Emerging Common Variants with Phenotype Correlation. Hum Mutat. doi: 10.1002/humu.22898

27. Kanagawa M, Kobayashi K, Tajiri M, Manya H, Kuga A, Yamaguchi Y (2016) Identification of a Post-translational Modification with Ribitol-Phosphate and Its Defect in Muscular Dystrophy. Cell Rep 14:2209–2223. doi: 10.1016/j.celrep.2016.02.017

28. Kurosawa H (2007) Methods for inducing embryoid body formation: in vitro differentiation system of embryonic stem cells. J Biosci Bioeng 103:389–398. doi: 10.1263/jbb.103.389

29. Li S, Edgar D, Fässler R, Wadsworth W, Yurchenco PD (2003) The role of laminin in embryonic cell polarization and tissue organization. Dev Cell 4:613–624. doi: 10.1016/S1534-5807(03)00128-X

30. Li S, Harrison D, Carbonetto S, Fässler R, Smyth N, Edgar D, Yurchenco PD (2002) Matrix assembly, regulation, and survival functions of laminin and its receptors in embryonic stem cell differentiation. J Cell Biol 157:1279–1290. doi: 10.1083/jcb.200203073

31. Li S, Qi Y, McKee K, Liu J, Hsu J, Yurchenco PD (2017) Integrin and dystroglycan compensate each other to mediate laminin-dependent basement membrane assembly and epiblast polarization. Matrix Biol 57:272–284. doi: 10.1016/j.matbio.2016.07.005

32. Lippmann ES, Estevez-Silva MC, Ashton RS (2014) Defined human pluripotent stem cell culture enables highly efficient neuroepithelium derivation without small molecule inhibitors. Stem Cells 32:1032–1042. doi: 10.1002/stem.1622

33. Meilleur KG, Zukosky K, Medne L, Fequiere P, Powell-Hamilton N, Winder TL, Alsaman A, El-Hattab AW, Dastgir J, Hu Y, Donkervoort S, Golden JA, Eagle R, Finkel R, Scavina M, Hood IC, Rorke-Adams LB, Bönnemann CG (2014) Clinical, Pathologic, and Mutational Spectrum of Dystroglycanopathy Caused by LARGE Mutations. J Neuropathol Exp Neurol 73:425–441

34. Michele DE, Barresi R, Kanagawa M, Saito F, Cohn RD, Satz JS, Dollar J, Nishino I, Kelley RI, Somer H, Straub V, Mathews KD, Moore SA, Campbell KP (2002) Post-translational disruption of dystroglycan-ligand interactions in congenital muscular dystrophies. Nature 418:417–421. doi: 10.1038/nature00837

35. Moore SA, Saito F, Chen J, Michele DE, Henry MD, Messing A, Cohn RD, Ross-Barta SE, Westra S, Williamson RA, Hoshi T, Campbell KP (2002) Deletion of brain dystroglycan recapitulates aspects of congenital muscular dystrophy. Nature 418:422–425. doi: 10.1038/nature00838

36. Morrissey MA, Sherwood DR (2015) An active role for basement membrane assembly and modification in tissue sculpting. J Cell Sci 128:1661–1668. doi: 10.1242/jcs.168021

37. Nakagawa N, Yagi H, Kato K, Takematsu H, Oka S (2015) Ectopic clustering of Cajal–Retzius and subplate cells is an initial pathological feature in Pomgnt2-knockout mice, a model of dystroglycanopathy. Sci Rep 5:11163. doi: 10.1038/srep11163

38. Nickolls AR, Bönnemann CG (2018) The roles of dystroglycan in the nervous system: insights from animal models of muscular dystrophy. Dis Model Mech 11:dmm035931. doi: 10.1242/dmm.035931

39. Peng HB, Ali A, Daggett DF, Rauvala H, Hassell JR, Smalheiser NR (1998) The relationship between perlecan and dystroglycan and its implication in the formation of the neuromuscular junction. Cell Adhes Commun 5:475–489. doi: 10.3109/15419069809005605

40. Praissman JL, Willer T, Sheikh MO, Toi A, Chitayat D, Lin Y-Y, Lee H, Stalnaker SH, Wang S, Prabhakar PK, Nelson SF, Stemple DL, Moore SA, Moremen KW, Campbell KP, Wells L (2016) The functional O-mannose glycan on alpha-dystroglycan contains a phospho-ribitol primed for matriglycan addition. Elife 5:e14473. doi: 10.7554/eLife.14473

41. Sanes JR (1982) Laminin, fibronectin, and collagen in synaptic and extrasynaptic portions of muscle fiber basement membrane. J Cell Biol 93:442–451. doi: 10.1083/jcb.93.2.442

42. Schittny JC, Timpl R, Engel J (1988) High resolution immunoelectron microscopic localization of functional domains of laminin, nidogen, and heparan sulfate proteoglycan in epithelial basement membrane of mouse cornea reveals different topological orientations. J Cell Biol 107:1599–1610. doi: 10.1083/jcb.107.4.1599

43. Sudo A, Kanagawa M, Kondo M, Chiyomi I, Kobayashi K, Endo M, Minami Y, Aiba A, Toda T (2018) Temporal requirement of dystroglycan glycosylation during brain development and rescue of severe cortical dysplasia via gene delivery in the fetal stage. Hum Mol Genet 27:1174–1185. doi: 10.1093/hmg/ddx143

44. Talts JF, Andac Z, Göhring W, Brancaccio A, Timpl R (1999) Binding of the G domains of laminin α1 and α2 chains and perlecan to heparin, sulfatides, α-dystroglycan and several extracellular matrix proteins. EMBO J 18:863–870. doi: 10.1093/emboj/18.4.863

45. Ungrin MD, Joshi C, Nica A, Bauwens C, Zandstra PW (2008) Reproducible, ultra high-throughput formation of multicellular organization from single cell suspension-derived human embryonic stem cell aggregates. PLoS One 3. doi: 10.1371/journal.pone.0001565

46. Vajsar J, Ackerley C, Chitayat D, Becker LE (2000) Basal lamina abnormality in the skeletal muscle of Walker-Warburg syndrome. Pediatr Neurol 22:139–143. doi: 10.1016/S0887-8994(99)00129-0

47. Winograd-Katz SE, Fässler R, Geiger B, Legate KR (2014) The integrin adhesome: From genes and proteins to human disease. Nat Rev Mol Cell Biol 15:273–288. doi: 10.1038/nrm3769

48. Yamamoto T, Toyoda C, Kobayashi M, Kondo E, Saito K, Osawa M (1997) Pial-glial barrier abnormalities in fetuses with Fukuyama congenital muscular dystrophy. Brain Dev 19:35–42. doi: 10.1016/s0387-7604(96)00056-3

49. Yoshida-Moriguchi T, Campbell KP (2015) Matriglycan: a novel polysaccharide that links dystroglycan to the basement membrane. Glycobiology 25:702–713. doi: 10.1093/glycob/cwv021

50. Yoshioka M, Kobayashi K, Toda T (2017) Novel FKRP mutations in a Japanese MDC1C sibship clinically diagnosed with Fukuyama congenital muscular dystrophy. Brain Dev 39:869–872. doi: 10.1016/j.braindev.2017.05.013

51. Yurchenco PD (2011) Basement Membranes: Cell Scaffoldings and Signaling Platforms. Cold Spring Harb Perspect Biol 3:a004911. doi: 10.1101/cshperspect.a004911

52. Zhang P, Yang Y, Candiello J, Thorn TL, Gray N, Halfter WM, Hu H (2013) Biochemical and biophysical changes underlie the mechanisms of basement membrane disruptions in a mouse model of dystroglycanopathy. Matrix Biol 32:196–207. doi: 10.1016/j.matbio.2013.02.002

